# Optimizing Proteomics Data Differential Expression Analysis via High-Performing Rules and Ensemble Inference

**DOI:** 10.1101/2023.06.26.546625

**Authors:** Hui Peng, He Wang, Weijia Kong, Jinyan Li, Wilson Wen Bin Goh

**Affiliations:** Lee Kong Chian School of Medicine, Nanyang Technological University, Singapore; School of Biological Sciences, Nanyang Technological University, Singapore; Shenzhen Institute of Advanced Technology, Chinese Academy of Sciences, China; Center for Biomedical Informatics, Nanyang Technological University, Singapore

## Abstract

In the process of identifying phenotype-specific or differentially expressed proteins from proteomic data, a standard workflow consists of five key steps: raw data quantification, expression matrix construction, matrix normalization, missing data imputation, and differential expression analysis. However, due to the availability of multiple options at each step, selecting ad hoc combinations of options can result in suboptimal analysis. To address this, we conducted an extensive study involving 10,808 experiments to compare the performance of exhaustive option combinations for each step across 12 gold standard spike-in datasets and three quantification platforms: FragPipe, MaxQuant, and DIA-NN. By employing frequent pattern mining techniques on the data from these experiments, we discovered high-performing rules for selecting optimal workflows. These rules included avoiding normalization, utilizing MinProb for missing value imputation, and employing limma for differential expression analysis. We found that workflow performances were predictable and could be accurately categorized using average F1 scores and Matthew’s correlation coefficients, both exceeding 0.79 in 10-fold cross-validations. Furthermore, by integrating the top-ranked workflows through ensemble inference, we not only improved the accuracy of differential expression analysis (e.g., achieving a 1-5% gain under five performance metrics for FragPipe), but also enhanced the workflow’s ability to aggregate proteomic information across various levels, including peptide and protein level intensities and spectral counts, providing a comprehensive perspective on the data. Overall, our study highlights the importance of selecting optimal workflow combinations and demonstrates the benefits of ensemble inference in improving both the accuracy and comprehensiveness of proteomic data analysis.

## Introduction

Proteomic data analysis workflows for differential expression analysis (DEA) usually comprise five key steps: quantifying raw data, constructing an expression matrix, normalizing the matrix, imputing missing data, and finally, conducting differential expression analysis by means of a statistical method. DEA workflows are crucial for accurately detecting phenotype-specific proteins potentially valuable in biomedical applications such as biomarker and drug target discovery ^[1, 2]^. Obtaining a good analysis outcome depends on identifying the most appropriate choice at every step to produce an optimal workflow (see **Figure 1**). However, as there are many options for each step in the DEA workflow, determining the optimal combination for a given data is challenging.

**Figure 1.**
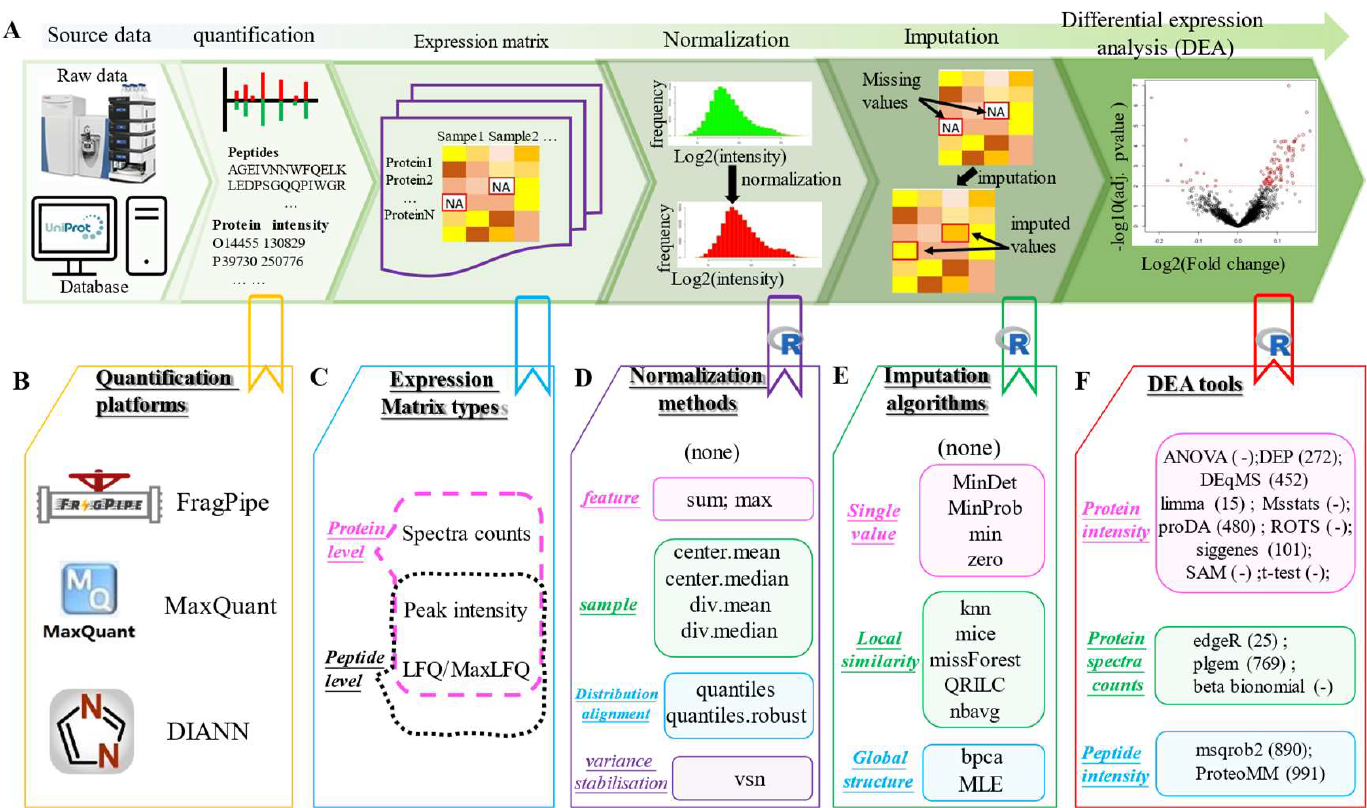
Workflow for DEA on proteomics data and available methods/tools for each step in the workflow. Panel **A** shows the five steps of a DEA workflow; **B**. peptide identification and quantification for protein assembling and quantitative analysis at a given analysis platform, e.g., FragPipe ^[13]^ for DDA or DIA-NN ^[14]^ for DIA; **C**. selection of quantification results organised as an expression matrix containing spectral counts or protein intensities; **D** normalization of raw expression matrix to reduce systemic bias, e.g., by center.mean or by vsn ^[15]^; and **E**. Imputation of missing values using methods such as missForest ^[16]^; and **F**. Selection of suitable DEA tool, e.g., ROTS ^[6]^ to identify truly differentially expressed proteins (DEPs) from non-DEPs. The numbers in the brackets show the user download frequencies which indicate their popularity.

There have been attempts to identify combinations of the steps for optimal DEA workflows but each of these does have limitations. Langley et al. ^[3]^ evaluated 7 DEA tools based on data of spectral count but did not consider the influence by other preprocessing steps such as normalization and missing value imputation (MVI) or intensity-based quantifications. Ramus et al. ^[4]]^ benchmarked 8 DDA (data-dependent acquisition) workflows on their in-house yeast dataset integrating different database search, protein assembly and validation, intensity-based quantitation, and two DEA tools; however, the impact of the data preprocessing methods was not considered (on yeast1819, see **Table 1**).

**Table 1.** Datasets used for workflow benchmarking.

| dataset | ID | acquisition | Sample | size | replicates | Contrasts |
| --- | --- | --- | --- | --- | --- | --- |
| yeast2099 | PXD002099* | DDA | yeast+48UPS1 | 5 | 3 | 10 |
| yeast1819 | PXD001819 | DDA | yeast+48UPS1 | 9 | 3 | 36 |
| human_ecoli_DDA | PXD018408 | DDA | Ecoli+HEK293 | 2 | 8 | 1 |
| human_ecoli_DIA | PXD018408 | DIA | Ecoli+HEK293 | 2 | 8 | 1 |
| CPTAC-LTQ86 | PDC000006** | DDA | yeast+48UPS1 | 5 | 3 | 10 |
| CPTAC-LTQO65 | PDC000006 | DDA | yeast+48UPS1 | 5 | 3 | 10 |
| CPTAC-LTQP65 | PDC000006 | DDA | yeast+48UPS1 | 5 | 3 | 10 |
| CPTAC-LTQW56 | PDC000006 | DDA | yeast+48UPS1 | 5 | 3 | 10 |
| RD139_Narrow | PXD026600 | DIA | Ecoli+48UPS1 | 8 | 3 | 28 |
| RD139_Wide | PXD026600 | DIA | Ecoli+48UPS1 | 8 | 3 | 28 |
| RD139_Overlap | PXD026600 | DIA | Ecoli+48UPS1 | 8 | 3 | 28 |
| RD139_Variable | PXD026600 | DIA | Ecoli+48UPS1 | 8 | 3 | 28 |

\* ProteomeXchange ID (http://proteomecentral.proteomexchange.org/cgi/GetDataset); \*\* Proteomic Data Commons Study Identifier (https://proteomic.datacommons.cancer.gov/pdc/)

Välikangas et al. ^[5]^ integrated several proteomics data processing (identification and quantification) software tools and evaluated nine MVI algorithms, but only one DEA tool ROTS ^[6]^ was compared, making their approach more suitable just for evaluating the impact of MVI. Fröhlich et al. ^[7]^ focused on evaluating DIA (data-independent acquisition) data processing tools, combining various DIA data processing tools, 4 normalization methods, and 7 DEA tools, but did not consider imputation algorithms. Lin et al. ^[8]^ compared DEA performance using tools originally designed for gene expression data and considered MVI and quantification tools, but not normalization methods. Sticker et al. ^[9]^ compared 7 DEA tools on spike-in datasets but ignored preprocessing methods. Dowell et al. ^[10]^ generated spike-in datasets to benchmark combinations of acquisition methods, replicate number, statistical approach, and FDR (False Discovery Rate) corrections ^[11]^, but only two DEA tools and one concentration contrast were tested, where the impact of preprocessing steps was not discussed. Suomi et al. ^[12]^ contrasted their DEA tool with 3 others but neglected data preprocessing.

Despite these studies, the impact of data preprocessing steps on DEA, especially the interdependence or synergies among these preprocessing steps, e.g., normalization and MVI algorithms together or not, remains poorly understood.

This work advances the study of DEA workflows for label-free proteomics data. We adopt a comprehensive approach by considering all possible workflows combining the choices available at each step for the quantification platforms, expression matrix types, normalization methods, imputation algorithms, and DEA tools. We evaluate these workflows across several publicly available spike-in datasets acquired in both DDA and DIA modes. Our leave-one-project-out cross-validations (using one proteomics project data as test data while the remaining project data are used for benchmarking to rank the workflow performances on the test data) confirm excellent generalizability of our benchmarking. We also applied a 10-fold cross validation to confirm whether a workflow’s performance level, e.g., high-performing or low-performing, is predictable from the options at each workflow step, e.g., normalization or not. Amongst the steps, we found that the choice of normalization methods and DEA statistical methods exert greater influence on performance over other steps. We also observed some interesting trends and frequent patterns amongst high-performing workflows, e.g., high-performing workflows prefer no normalization and incline MinProb (probabilistic minimum) for imputation while eschewing simple statistical tools (e.g., ANOVA ^[17]^, SAM ^[18]^ and t-test ^[19]^) for DEA. By studying the impact of choices at each step of a workflow on DEA performance, we provide a unique resource to guide workflow selection on new datasets. In addition, we introduce a powerful ensemble inference strategy that integrates DEA results from individual top-performing workflows to further improve DEA performance. This strategy improves DEA performance by 1∼5% (on 5 different kinds of performance metrics) for FragPipe-based or 1∼4% for maxquant-based DDA data ^[20]^ (non-G-mean ^[21]^ metrics), and 2∼4% for DIA-NN-based DIA data. While we also found that spectral count may not work as well as intensity (at both protein and peptide level) in DEA, the combination of these multiple proteomic data layers provides complementary information that enhances DEA outcomes. We therefore note that an ensemble approach can help maximize available information in a single dataset by harnessing its multiple perspectives.

## Results

### Details of proteomic datasets

We amassed 7 DDA datasets and 5 DIA datasets from 5 proteomics projects for assessing DEA workflows. **Table 1** presents information on acquisition type, sample size, replicates, and contrasts (a contrast refers to the expression level comparison of two groups of proteins acquired from two different samples. Thus, the contrast number in a dataset equals the number of unique sample pairs in it).

Altogether, we considered 40 DDA datasets (1 yeast2099 ^[22]^ + 1 yeast1819 ^[4]^ + 34 human_ecoli_DDA ^[10]^ + 4 CPTAC ^[23]^, the 34 human_ecoli_DDA datasets were generated by bootstrapping, more details see **Supplementary results**) and 38 DIA datasets (4 RD139 ^[24]^ + 34 human_ecoli_DIA ^[10]^). For a given spike-in dataset, e.g., the CPTAC_LTQ86, 48 UPS1 (Universal Proteomics Standard 1) proteins with 5 concentrations were added to the same yeast background proteins. 10 contrasts can be made if we choose samples with 2 out of the 5 different concentrations (*C*^2^ = 10) for comparison to retrieve differentially expressed proteins. Thus, there are 120 (10+36+34+10 * 4) contrasts for DDA datasets and 146 (28 * 4+34) contrasts for DIA datasets. Each contrast was evaluated by different workflows.

We used 3 popular and freely accessible quantification software platforms for quantitative analysis: Fragpipe ^[13]^ and maxquant ^[20]^ for DDA data and DIA-NN ^[14]^ for DIA data. As each of these platforms use different database search, peptide identification, protein assembly and quantification algorithms, we may get drastically different answers given the same dataset. Moreover, the compatibility of the data with the workflow, e.g., no spectral count data is available from DIA-NN, is another factor worth considering.

### Optimal workflows are identifiable through benchmarking and are platform-specific

Candidate workflows were evaluated on 5 metrics: partial area under receiver operator characteristic curves (pAUC) ^[25]^ with false-positive rate (FPR) thresholds of 0.01, 0.05, 0.1 (pAUC(0.01), pAUC(0.05), pAUC(0.1)), normalized Matthew’s correlation coefficient (nMCC) ^[26]^ and geometric mean of specificity and recall (G-mean) ^[21]^ (See **Method** for details). **Figure 2A** shows the distributions of the five performance-metrics obtained by FragPipe-related workflows on different datasets. Some workflows produce very good performance whereas others lead towards extremely poor results. This shows that optimal workflow selection is important.

**Figure 2.**
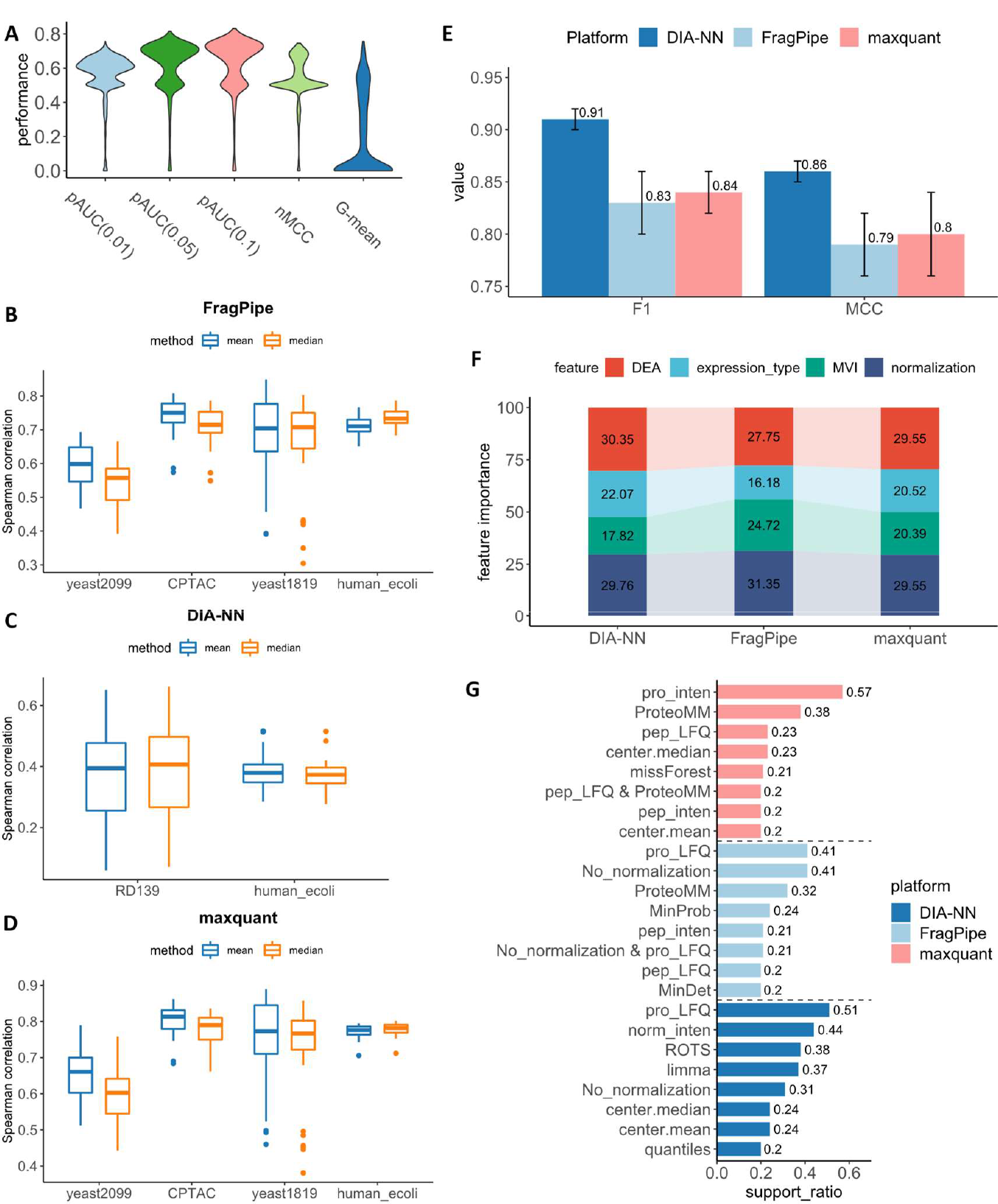
Performance distributions of workflows, leave-one-project-out cross-validation test results and data mining-based analysis on high-performing workflows. Panel **A** shows the performance distribution of FragPipe-based workflows. The workflows are labeled as “H”, “RH”, “RL” and “L” if their average performances are ranked at top 5% (there are 151 “H” workflows), between 5% and 25% (600 “RH” workflows), between 25% and 50% (751 “RL” workflows) and below 50% (1502 workflows). Panel **B, C** and **D** present leave-one-project-out cross-validation test results for evaluation of performances of applying benchmarking results to guide workflow selection for new coming datasets. Panel **E** gives 10-fold cross validation performances of workflow classification with CatBoost classifiers. Panel **F** shows feature importance of CatBoost classifiers in workflow classification. Panel **G** lists the found frequent patterns with support ratios bigger than 0.2 from “H” workflows.

We use the mean or median performance of a workflow across multiple datasets to establish its overall performance ranking. Furthermore, a workflow’s final rank is given by averaging its five independent ranks on 5 metrics (see **Method**). We use leave-one-project-out cross-validation (LOPOCV, a type of leave-one-out cross-validation, where in each round, the dataset from the same proteomics project is left for testing and remaining data are used for benchmarking, see **Method** for more details) to test whether our benchmarking results (final ranks of workflows) can benefit recommendations of workflows for newcoming datasets (generalizability). We use the Spearman correlation coefficient to measure consistency between workflow ranks obtained by our benchmarking and a new dataset’s true performance.

In **Figure 2B-D**, we show the LOPOCV results tested on FragPipe, maxquant and DIA-NN workflows respectively. From **Figure 2B** we can see, testing on the CPTAC project (total 40 tests), average spearman correlation of 0.74 is obtained when using mean performance of multiple datasets to rank FragPipe workflows. If median performance is used, an average spearman correlation of 0.71 is obtained. For projects yeast2099 (10 tests), yeast1819 (36 tests) and human_ecoli (34 tests), their average spearman correlations are 0.59(0.54), 0.68(0.66) and 0.71(0.74) for mean(median) performance-based benchmarking. For maxquant workflows (**Figure 2D**), average spearman correlations of 0.65(0.60), 0.80(0.78), 0.74(0.72) and 0.77(0.78) are achieved for mean(median) performance-based workflow benchmarking. In **Figure 2C**, we show the evaluation of DIA-NN workflow benchmarking. For the two DIA projects RD139 (112 tests) and human_ecoli (34 tests), much worse average spearman correlations of 0.37(0.38) and 0.38(0.38) are obtained. This may be due to less project data (only 1 project) being used for workflow benchmarking, resulting in poorer generalizability. This has been proved by our benchmarking data size affection tests, where fewer data always results in worse LOPOCV average Spearman correlations (see **Supplementary results and Supplementary file 1**). For example, when testing on CPTAC project data generated by FragPipe, if we randomly selected 1 (or 2) project data for workflow benchmarking with mean performance, then the average Spearman correlations will drop to 0.67 (or 0.72). In conclusion, our benchmarking ranks are highly positively correlated to true workflow performances (overall average correlations > 0.7 for FragPipe and maxquant), which makes it powerful for guiding optimal workflow selection. In addition, mean performance works a little better than median performance for workflow benchmarking in most LOPOCV tests. Mean performance is used for workflow benchmarking and in the following analysis.

Top ranked workflows for FragPipe, maxquant and DIA-NN appear rather varied in the four steps (i.e., platform-wised ranking) following specification of the quantification platform. For FragPipe, the best workflow comprises expression matrix type of peptide MaxLFQ ^[27]^ intensity, max normalization, MinDet (deterministic minimum) MVI and ProteoMM ^[28]^ for DEA. This workflow has a mean pAUC(0.01) score of 0.76, a mean pAUC(0.05) score of 0.82, a mean pAUC(0.1) score of 0.83, a mean nMCC score of 0.73 and a mean G-mean of 0.76. For maxquant, the top workflow comprises peptide peak intensity as input expression matrix, vsn ^[15]^ for normalization, imputation via MinDet and ProteoMM for DEA; its performance under the five-mean metrics is 0.73, 0.78, 0.80, 0.72 and 0.67 (in the same order as FragPipe). For DIA-NN, the best workflow comprises protein ‘norm’ intensity ^[27]^, no normalization (None), QRILC ^[29]^ for imputation and ROTS for DEA; and its mean performances are pAUC(0.01) = 0.88, pAUC(0.05) = 0.92, pAUC(0.1) = 0.93, nMCC = 0.88 and G-mean = 0.81.

We observed large performance gaps between top- and bottom-ranked workflows. For instance, among the 4800 DIA-NN workflows, the 2701^st^-ranked workflow “protein ‘norm’ intensity + ‘quantiles.robust’ normalization + QRILC MVI+ ROTS” has the five mean metrics scores of 0.58, 0.65, 0.66, 0.55 and 0.19. This is quite dramatic, showing that changing just one step, in this case, the normalization method, can transform a top-ranked workflow into a bottom-ranked one. This phenomenon also occurs in workflows pertinent to other platforms (see **Supplementary file 2**).

### Extracting frequent patterns from high-performing workflows

We use machine learning to identify common traits useful for predicting high-performing workflows. For each platform, we encoded every workflow as a feature vector by regarding every option in a step as a categorical feature value. We assigned each workflow a performance level such as high (“H”), relatively high (“RH”), relatively low (“RL”) or low (“L”), if its rank is within top 5%, between top 5% and top 25%, between top 25% and 50%, and the remaining 50% respectively (see **Supplementary file 3**). We used these workflow feature vectors and their labels to train a CatBoost classifier ^[30]^ followed by 10-fold cross validation for performance evaluation (**Figure 2E**). We found that the performance level of a workflow is predictable because the resultant F1 or MCC values are high with an average of 0.79 and above. In addition, we also found that model performance is influenced more by the choice of normalization and DEA tool than expression matrix type and MVI algorithms (**Figure 2F**; see **Supplementary file 3** for more details).

We focused on the “H” workflows, so that we may discover rules or patterns associated with high performing workflows. We used the Frequent Patten Growth (FP-growth) algorithm ^[31]^ to identify frequent selection patterns in “H” labeled workflows (we repeated analyses on “L” workflows for comparison). Altogether, we studied 151, 151 and 240 “H” workflows for FragPipe, maxquant and DIA-NN respectively. Frequent patterns with a support ratio (SR, defined as the fraction of the total workflows containing the pattern) greater than 20% are depicted in **Figure 2G (**frequent patterns with SR>=10% but SR<=20% are listed in **Supplementary file 4)**.

We identified some common frequent selection patterns for “H” workflows across three quantification platforms. Normalization methods “center.mean”, “center.median” and “None” (no normalization) are frequently selected for all three platforms with support ratio bigger than 10% (see **Supplementary file 4**). For FragPipe and DIA-NN, “None” is the most frequent choice for normalization method selection (SR=0.41 for FragPipe and SR=0.31 for DIA-NN). In comparison, normalization methods of “sum”, “max”, “div.mean”, “div.median”, “quantiles.robust” are non-favorable (as they are frequently selected in “L” workflows). For MVI algorithms, “MinProb” and “MinDet” are frequently selected to impute missing values by “H” workflows for the three platforms ^[8]^. Similarly, “H” workflows favored limma ^[32]^ as the DEA of choice (with SRs of 16%, 13%, 37%) whereas simple statistical tools, e.g., ANOVA, SAM and t-test are enriched in low performing “L” workflows ^[33]^.

Some of the high-performing single step choices reported above are unsurprising. However, our approach can also identify interesting synergies (or “avoidances”). For example, no normalization is coupled with protein MaxLFQ intensity in both FragPipe (SR=21%) and DIA-NN (SR=17%) “H” workflows. No normalization is not only coupled with protein peak intensity in FragPipe and maxquant “H” workflows but also coupled with “norm” intensity in DIA-NN “H” workflows (SRs>10%). Both peptide peak intensity and peptide MaxLFQ intensity are coupled with ProteoMM in FragPipe and maxquant “H” workflows (SRs>15%). In DIA-NN “H” workflows, MaxLFQ intensity is coupled with limma (18%) and ROTS (16%). The pair (protein MaxLFQ intensity, limma) also frequently appears in FragPipe “H” workflows (12%). The pairs (protein peak intensity, limma) and (protein peak intensity, DEqMS ^[34]^) are selected in maxquant “H” workflows with SR=13% and SR=17% respectively. In comparison, no coupled pattern is identifiable in “L” workflows (see **Supplementary file 4** for more details). In addition, we identified platform-specific frequent patterns. For example, protein MaxLFQ intensity (SR=41%) and no normalization (SR=41%) for FragPipe, protein peak intensity for maxquant (SR=57%) and MaxLFQ intensity for DIA-NN (SR=51%) (**Figure 2G**). These selections are highly recommended.

Some workflow selection guidelines are derived from these frequent patterns. First, the platform-specific frequent patterns should always be given priority for inclusion in workflow. Suppose we need to construct a FragPipe workflow, we should opt to use protein MaxLFQ intensity together with no normalization as these options by themselves are not only FragPipe-specific frequent patterns but also constitute a synergistic combination (SR=21%) in FragPipe “H” workflows. Since we are using protein MaxLFQ, we opt for limma as a DEA selection as it is positively coupled with protein MaxLFQ intensity (SR=12%). We suggest also using these selections to help guide an appropriate MVI algorithm, e.g., MinProb (if chosen, the workflow is ranked at 11^th^) or MinDet (ranking at 12^th^) to guarantee a good performance.

Similarly, for maxquant, we recommend choosing protein peak intensity (maxquant-specific) and coupling this selection with DEqMS (or limma) for DEA and “center.median”, “center.mean” or “None” are favored for normalization (high SRs or synergy). Finally, an MVI method, e.g., MinProb, MinDet or missForest (high SRs) can be chosen to achieve a higher performance (of the 18 possible workflows, only 2 of them are ranked lower than 80^th^ and the worst one is ranked at 143^rd^).

For DIA-NN, MaxLFQ intensity is recommended, and this can be coupled with limma or ROTS (synergy) as the DEA tool. There are several good options for normalization: no normalization (synergy), “center.mean” and “center.median” (high SRs). And MinProb or MinDect (high SRs) are good MVI selections.

### Comparing expression matrix types, imputation, normalization, and DEA tools

We compared options in each workflow step (see **Method**). Only the performance indicator pAUC(0.01) was used for comparisons though 5 different indicators were selected for final workflow ranking (see **Method**). This is because pAUC(0.01)-based ranking of workflows always achieves the highest correlations with the final average rankings for all the three quantification platforms (Pearson r=0.88, Spearman R=0.89 for FragPipe; r=0.89 and R=0.90 for maxquant; and r=0.87 and R=0.88 for DIA-NN, see **Supplementary file 5**). Comparison results are shown in **Figure 3**.

**Figure 3.**
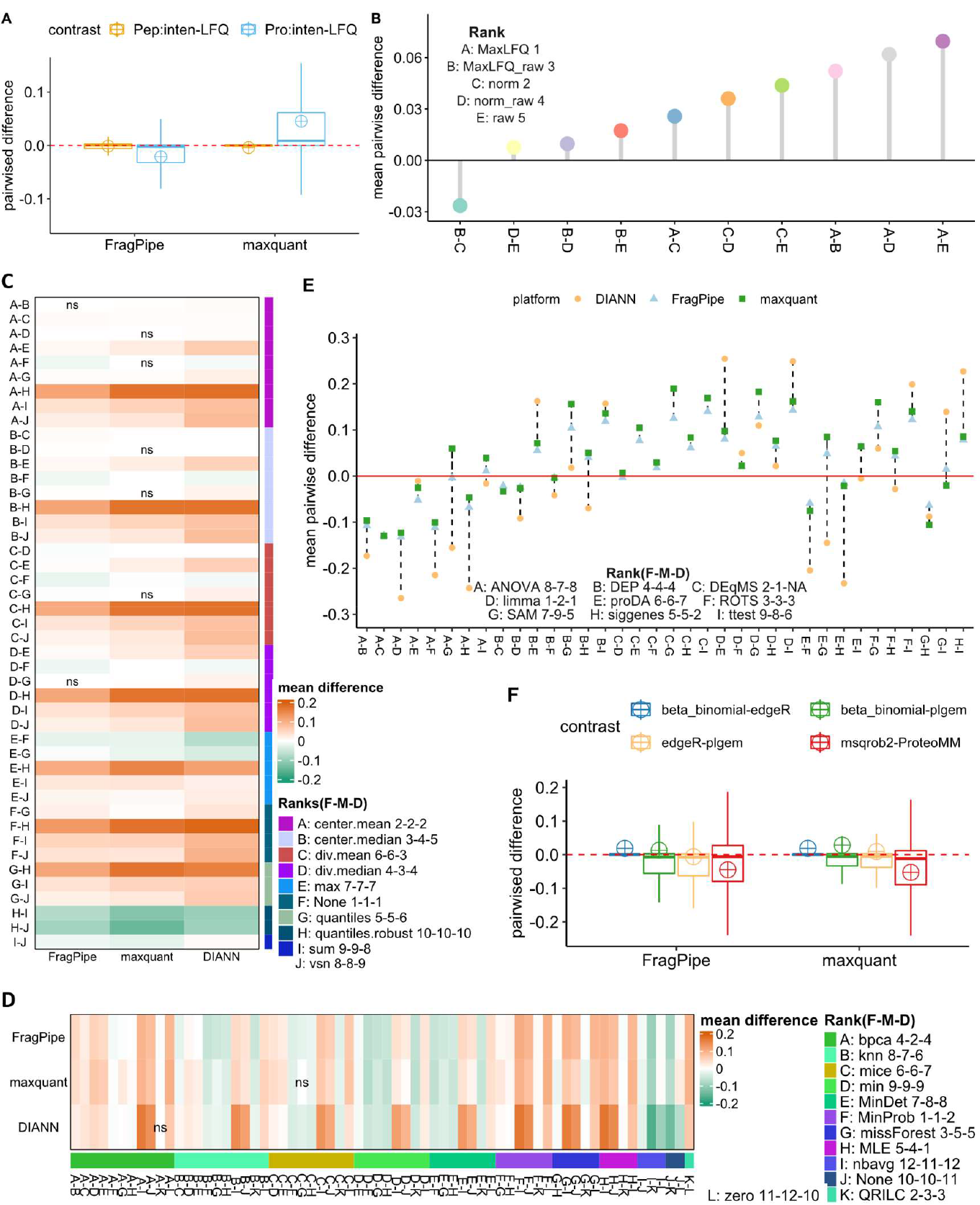
The impact of expression matrix type, normalization, imputation and DEA tool on differential expression analysis performance. **Panel A** shows the pairwise comparison of expression matrix types of FragPipe and maxquant. **B** compares five expression matrix types from DIA-NN. **C** gives the comparison results of various normalization methods and their rankings. **D** presents the median pairwise differences of different imputation algorithms and annotates their final rankings. **E** and **F** describe the performance differences of popular DEA tools under different quantification platforms and using different expression matrixes. Here, F-M-D represents FragPipe-maxquant-DIA-NN. Markers show mean performances. “ns” denotes “not significant” defined by the t-test p-value>0.05.

There is little performance difference between peptide peak intensity and peptide MaxLFQ intensity in both FragPipe (mean=-0.001, paired t-test p<1e-5) and maxquant (mean=-0.004, paired t-test p<1e-5) based DEAs (see **Figure 3A**). At protein level, FragPipe-related workflows prefer MaxLFQ intensity as expression matrix type (mean=-0.022 paired t-test p<1e-5) while peak intensities work a little better in maxquant-related workflows (mean=0.046, paired t-test<1e-5). For DIA data, use of MaxLFQ intensities is favorable (ranking 1^st^ according to pairwise comparisons in **Figure 3B**). This result is consistent with our frequent pattern mining results that in high-performing workflows, protein MaxLFQ intensity obtains the highest SRs, thus should be selected with priorities.

For normalization methods, the no normalization (F:None) is always ranked 1^st^, followed by center.mean (A) and center.median (B) where small decreases were obtained (within 0.01 shown with light colors in heatmap of **Figure 3C**, also see **Supplementary file 6**). However, for some normalization methods e.g., quantiles.robust, the DEA performances decreased significantly (bigger than 0.05). Surprisingly, this suggests that normalization may not be a necessary (or appropriate) preprocessing step for DEA workflows. As center.mean and center.median were frequently found in high-performing workflows, we recommend using center.mean and center.median (or even no normalization). This suggestion is made purely from the benchmark and does not mean that data normalization should be avoided at all costs. It is possible there are situations which warrant the use of normalization techniques, but these should be carefully customized to the data and research question at hand.

When comparing imputation methods, most MVI methods produce better performances than no imputation (J:None) (**Figure 3D** and **Supplementary file 6**). Among them, MinProb always works well across three platforms. This is not surprising, as MinProb addresses missing-not-at-random (MNAR) missingness, which plagues proteomics data ^[35]^. Alternatively, QRILC, missForest and bpca ^[36]^ are also good selections with small performance differences comparing to MinProb (**Supplementary file 6**) and are ranked high (**Figure 3D**).

When comparing DEA tools, limma is always the best DEA tool for FragPipe and DIA-NN related workflows (**Figure 3E**). DEqMS, designed specifically for MS data, works well, and performs better than most tools adapted from elsewhere, e.g., ROTS and SAM (originally for microarray). For spectral count-based DEA, beta-binomial works slightly better while for peptide level DEA, ProteoMM outperforms msqrob2 ^[37]^ (**Supplementary file 6 and Figure 3F**). Though we cannot compare protein intensity, peptide intensity and spectral count-based tools in DDA data DEA directly, we find that trend-wise, spectral count-based workflows perform worse than peptide intensity and protein intensity-based workflows (no high-performance workflows involve spectral count). In comparison, about 3/5 of high-performing workflows are associated with protein intensity and the remaining 2/5 are correlated with peptide intensity (see **Supplementary file 3**).

### Ensemble inference by integrating top ranked workflows improves DEA performance

This work highlights the importance of considering multiple DEA workflows as they can provide complementary information and improve the overall coverage of differentially expressed proteins. Despite having similar performance scores, top-ranking workflows may not report the same set of differentially expressed proteins. Integrating results from multiple high-performing workflows can provide a more comprehensive view of the differential expression landscape and increase the confidence of the results. Indeed, Goeminne et. al. proved that uniting intensity-based and spectral counts-based analysis can improve DEA performances ^[38]^. Moreover, those false positives of differentially expressed proteins have the chance to be voted down by considering the outcomes of multiple DEA workflows. Additionally, the combination of results from different workflows (even if they use the same expression matrices) can increase the robustness of the results while reducing false positives/negatives. Hence, we conduct ensemble inference by integrating the outcomes from top-ranking workflows.

Our results show that ensemble inference strategies, ens_3inp, ens_2inp, and ens_TK (see **Method**) improve DEA performance by providing comprehensive and accurate estimation of differential expressions. **Figure 4A** shows the pAUC(0.01) score distributions obtained by the best spectral count-based workflow (spec_T1), the best protein intensity-based workflow (pro_T1), the best peptide intensity-based workflow (pep_T1), the best ensemble inference with ens_3inp, the best ensemble inference with ens_2inp and the best ensemble inference with ens_TK testing on different DDA datasets and using FragPipe as the quantification platform. Here, ens_3inp integrates spec_T1, pro_T1 and pep_T1 by using hurdle (see **Method**) to calculate integrated p-values, ens_2inp integrates pro_T1 and pep_T1 with hurdle, while ens_TK integrates top 7 workflows (K = 7), and minimum p-values was used.

**Figure 4.**
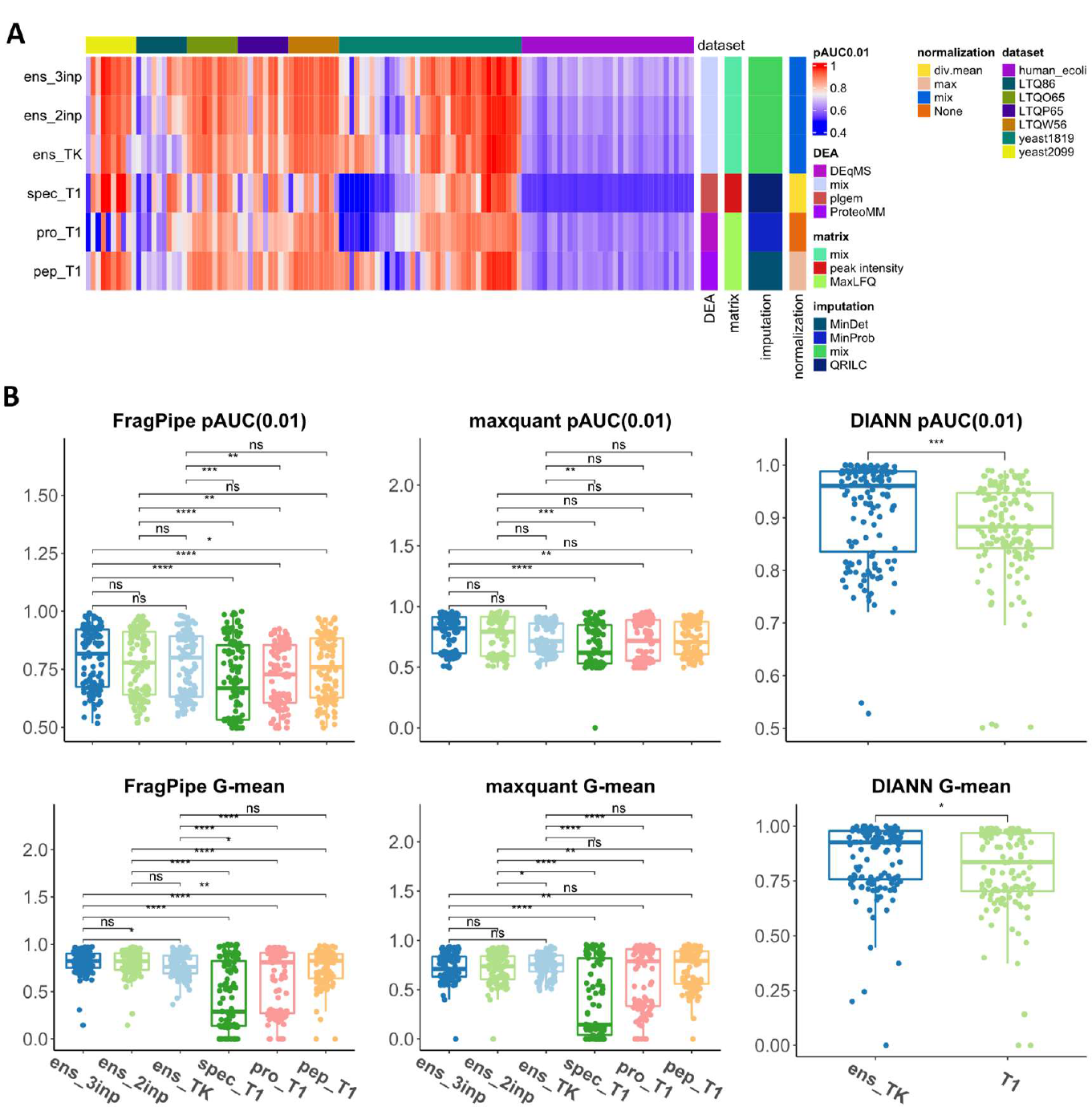
Benchmarking and ensemble inference results. **Panel A** shows an example of pAUC(0.01) value profile achieved by FragPipe related top-ranking workflows or ensemble inferences tested on different datasets. **B** compares the ensemble inferences and top-ranking single workflows under three quantification platforms.

From **Figure 4A**, we can see both ens_3inp, ens_2inp and ens_TK reduce blue cells compared to spec_T1, pro_T1 and pep_T1 especially on those left-panel datasets, which means higher pAUC(0.01) scores were obtained. To clearly compare these local optimal workflows (ranking first for a specific expression matrix type or ensemble strategy), we analyzed their performance distributions (**Figure 4B**).

**Figure 4B** compares results from different local optimal workflows with one representative pAUC based-performance indicator pAUC(0.01) and one classification-based performance indicator G-mean. FragPipe-related comparisons are shown in the first column followed by maxquant-related comparisons and then DIA-NN-related comparisons (only ens_TK and pro_T1). T-test derived p-values are drawn above the boxplots. When using FragPipe as the quantification platform, ens_3inp (ens_3inp with hurdle) significantly improves DEA performance by about 1∼5% of the five-performance metrics (their mean values are [0.80, 0.86, 0.87, 0.74, 0.81]) compared with the best overall ranking workflow. In comparison, ens_2inp obtained 0∼4% performance improvement and ens_TK (ens_T7 with min) outperformed other single workflows on most metrics (1∼2% increases except for nMCC(−2%)). For maxquant, ens_3inp, ens_2inp and ens_TK made modest gains on most metrics (see **Figure 4B** middle column, ens_3inp obtains 1∼4% performance gain for non-G-mean indicators). For DIA-NN, there was significant improvement by using ens_TK (K=20, min p (see **Method**)) especially when compared against pro_T1 (2∼4% performance gain).

From the first two columns of **Figure 4B** and the comparisons between the ens_3inp and ens_2inp (with/without spectral count, see our **Supplementary file 7**), we observe that while spectral count-based workflows perform worse than protein intensity or peptide intensity-based workflows (spectral count also frequently appears in FragPipe (SR=18%) and maxquant “L” (SR=19%) workflows), they may be integrated with protein intensity or peptide intensity-based workflows to achieve performance gains.

Though at most 5% improvement could be achieved, ensemble inference is highly recommended to find more accurate differential expressed proteins. For DDA data, ens_3inp (hurdle) always works the best compared against ens_3inp with other integration strategies, ens_2inp with different integration strategies, or compared against ens_TK (also see **Supplementary file 7**). Since it is easy to obtain protein intensities, peptide intensities, and spectral counts from FragPipe and maxquant, we recommend using ens_3inp (hurdle) with DDA data. For DIA data processed by DIA-NN, ens_TK with bigger K, e.g., 15, 20, we recommend using minimum p.

### Guidelines for selecting optimal workflows

By exhaustively testing workflow combinations, we make the following recommendations:

1. For DEA with FragPipe’s quantification outputs, selecting a workflow combining protein MaxLFQ intensity, no normalization and limma at first, then, a MVI algorithm of MinProb or MinDect may provide high performance.
2. For DDA data analyzed with maxquant, protein peak intensity with no normalization, center.mean or center.median for normalization and limma (or DEqMS) for DEA, together with an imputation algorithm of MinProb, MinDet or missForest can result in higher performances.
3. For DIA data processed by DIA-NN, workflows integrating protein MaxLFQ intensity, the DEA tool of ROTS or limma, any normalization method drawn from the pool of (no normalization, center.median, center.mean) and a MVI algorithm drawn from the pool of (MinProb, MinDet) may assure good DEA performance.
4. For DEA tools, if spectral count is used for quantification, then beta-binomial is better. If protein intensities are used, we recommend limma, DEqMS and ROTS for FragPipe and maxquant, and Limma, siggenes ^[39]^ and ROTS for DIA-NN. For peptide level, ProteoMM is better than msqrob2.

The above points provide some broad suggestions on optimal workflows. Below, we also provide some tools to help users identify the best workflow for their data:

1. Select the top-ranking workflows according to our benchmarking results. We provided a webserver OpDEA (http://www.ai4pro.tech:3838/OpDEA_test/) and a R package (https://github.com/PennHui2016/OpDEA) to guide which workflow should be used after specifying the quantification platform and/or the preferred expression matrix types.
2. Use the ensemble inference workflow of ens_inp3 (hurdle) to improve overall DEA performance for DDA data if protein intensities, peptide intensities and spectral counts are all available or ens_inp2 (hurdle) if protein intensities and peptide intensities are available. For DIA data, ens_TK (K=20, min p) is recommended for ensemble inference.
3. Our benchmarking results (see **Supplementary file 2**) can be used as references for users to generate their self-defined workflows, where they can check their workflows’ ranking in our results and then tune them for optimization.

## Discussion

### Our ranked workflows provide a knowledge resource for users to identify optimal tools

Our workflows test an extensive set of steps and their combinations (including expression matrix types, expression data preprocessing methods for normalization and imputation, and DEA tools) to make a total of 10,808. We are therefore able to provide optimal workflow selection guidelines. We also evaluated broadly across a set of 5 performance indicators and across 3 quantification platforms to ensure balanced perspectives. We are therefore confident that the webserver provided here is valuable.

Our study highlights the significant impact of missing data on DEA performance and thus, the importance of paying attention to MVI algorithm selection.

Furthermore, we comprehensively investigated the impact of tool selections in each important step of a complex workflow. Our findings showed that expression matrix types have minor effects on DEA performance, while normalization may even worsen it. On the other hand, selection of a good MVI algorithm and a compatible DEA tool is highly beneficial.

### Limitations of current work

We maintained the default parameters for each tool to maintain consistency in the evaluation process. However, we note that some parameters may need to be adjusted based on the specific properties of the data. For example, the proteomics data generated by different spectrometers may have different resolutions and require different search parameters, such as precursor mass tolerance or fragment mass tolerance. However, these database search parameters do not impact the fairness of comparisons among downstream DEA workflows as they start from the quantification results, not the raw data.

Some imputation or DEA tools have tuneable parameters, such as “k” for “knn” imputation ^[40]^ or the number of bootstrapping for ROTS ^[6]^. While adjusting these parameters can affect the final DEA performance, it also increases the time cost and may require additional labelled golden-standard data for validation. This is not feasible for analysing real-life data.

By using the default parameters, we ensure a stable and general evaluation of the tools working on randomly chosen datasets. The default parameters are the best-practice values optimized by the developers and can produce commonly accepted performance. Our benchmarking based on default settings provides a reliable and consistent evaluation of the tools.

## Conclusion

We benchmarked 10,808 DEA workflows on 5 performance indicators across 40 DDA datasets and 38 DIA datasets. The results provide a simple and straightforward recommendation of the best workflow for a DEA task and give insights into the impact of different choices in each step of the DEA workflow. Our findings indicate that the selection of a good MVI algorithm, normalization method and DEA tool is more crucial for DEA performance compared to the choice of expression matrix type. Furthermore, we demonstrate that ensemble inference strategies using top-ranking workflows can significantly improve DEA performance. Our benchmarking results provide stable and general evaluation of the tools’ performance on randomly chosen datasets, and the results can be used to guide optimization of DEA workflows.

## Method

### Quantitative analysis of mass spectrometry -based proteomics data

*DDA data quantification*. FragPipe v17.1 software was used with default quantification settings. MSFragger-3.4 ^[13]^ was adopted for database search (using default parameters), contaminants and decoy (sequence reverse) protein sequences were added to corresponding libraries, e.g., 48 human UPS1 proteins + reviewed UP000002311 Saccharomyces cerevisiae proteome from uniport database ^[41]^ for yeast2099, yeast1819 and CPTAC datasets or 48 human UPS1 proteins + reviewed UP000000625 Escherichia coli proteome for RD139 datasets or reviewed UP000000625 Escherichia coli proteome + reviewed UP000005640 human proteome for human_ecoli datasets. Percolator ^[42]^ and ProteinProphet ^[43]^ were used for peptide identification and protein inference. For quantification, IonQuant ^[44]^ with match between runs ^[45]^ and MaxLFQ ^[27]^ were used. The quantification results are stored in files containing identified peptides or proteins with their spectral counts or intensities and other annotation information. We extracted expression levels of identified peptides or proteins in the form of spectral counts or intensities and organized them as a matrix by setting proteins as rows and samples as columns. The cells hold the expression levels. This data structure is referred to as an expression matrix, and it holds varied expression information. In fact, five types of expression matrixes are obtainable from FragPipe including spectral counts, protein level peak intensities and MaxLFQ intensities calculated by MaxLFQ algorithm ^[27]^ and peptide level peak intensities and MaxLFQ intensities.

MaxQuant v2.0.3.1 was used as an alternative quantification platform for DDA data ^[20]^. Andromeda ^[46]^ was applied for database search. We kept most parameters default and used the same reference library as FragPipe. Similarly, match-between runs and MaxLFQ label-free quantification algorithm was used. Again, spectral count, protein level peak intensities and MaxLFQ intensities, and peptide level peak intensities and MaxLFQ intensities can be acquired for DEA. We note that maxquant returned errors when processing the CPTAC_LTQP65 dataset (**Table 1**), so only 110 contrasts were tested by maxquant -based DEA workflows.

#### DIA data quantification

For DIA data, DIA-NN v1.8 was used for quantification with default parameters and predicted libraries from corresponding databases ^[14]^. Match between runs was checked and we run DIA-NN under low RAM and high-speed mode. Five types of protein intensities can be obtained from its output files such as normalized intensities (‘norm’) extracted from protein group report file (‘report.pg_matrix.tsv’) and another expression matrixes obtained with diann R package (https://github.com/vdemichev/diann-rpackage) including non-normalized intensity (‘raw’), normalized intensities (‘norm_raw’) and MaxLFQ extracted with ‘diann_matrix’ function from main report file (‘report.tsv’) (‘MaxLFQ_raw’), and MaxLFQ intensities extracted with ‘diann_maxlfq’ function from main report file (‘MaxLFQ’).

### Benchmarking settings of expression matrix type, normalization, imputation and DEA tools

For DDA data quantified with FragPipe and maxquant, five types of expression matrixes are obtainable including spectral counts, protein level peak intensities and MaxLFQ intensities calculated by MaxLFQ algorithm ^[27]^, and peptide level peak intensities and MaxLFQ intensities. As for DIA quantification with DIA-NN, five types of protein intensities are obtainable including ‘norm’, ‘raw’, ‘norm_raw’, ‘MaxLFQ_raw’, and ‘MaxLFQ’ (see above section).

Two preprocessing procedures (normalization and imputation) are conducted (**Figure 1A**). Many methods have been proposed to normalize expression data and impute missing values. It’s impossible to evaluate every normalization and imputation method. And so, only popular and readily accessible ones were used.

For normalization methods, we used those found in the MSnbase v2.22.0 R package ^[47]^ including “sum”, “max”, “center.mean”, “center.median”, “div.mean”, “div.median”, “quantiles”, “quantiles.robust” and “vsn”(variance stabilization) ^[15]^ (we conclude them in **Figure 1D**). If no normalization is used, then we designate the normalization method as “None”.

For MVI methods, we also used those found in the MSnbase v2.22.0 R package such as “bpca” (Bayesian principal component analysis) ^[36]^, “knn” (k-nearest neighbors) ^[40]^, “QRILC” (quantile regression imputation of left-censored data) ^[29]^, “MLE” (maximum likelihood estimation), “MinDet” (deterministic minimum), “MinProb” (probabilistic minimum), “min”, “zero” and “nbavg” (neighbour averaging), and another two popular imputation methods such as mice (Multivariate imputation by chained equations) ^[48]^ and missForest (random forest) ^[16]^. Again, if no imputation is performed, we designate the imputation method as “None”.

We identified DEA tools across available implementations in Bioconductor v3.15 ^[49]^ and those we collected from published literature. From Bioconductor (Bioconductor - BiocViews: https://www.bioconductor.org/packages/release/BiocViews.html__DifferentialExpression), we found DEA tools designed specifically for proteomics data. We ranked them by popularity given their download numbers (shown in brackets). Our list includes DEP (272) ^[50]^, DEqMS (452) ^[34]^, plgem (769) ^[51]^, proDA (480) ^[52]^, msqrob2 (890) ^[37]^, ProteoMM (991) ^[28]^. Popular and commonly used DEA tools limma (15) ^[32]^, edgeR (25) ^[53]^ and siggenes (101) ^[39]^ are not originally designed for proteomics data but are also downloaded quite frequently and made it into our list. Other general tools such as ANOVA ^[17]^, t-test ^[18]^, beta_binomial ^[54]^, Msstats ^[55]^, SAM ^[18]^ and ROTS ^[6]^ are collected from literature ^[7]^ and ^[38]^ or related websites. Each DEA tool has a preferred expression matrix type: edgeR, plgem and beta_binomial are good with spectral counts; ProteoMM and msqrob2 are compatible with peptide level intensities; while other tools are either agnostic or work well directly with protein intensities. Some other DEA tools are more demanding: MSstats and DEqMS require additional information beyond protein intensities where MSstats require feature-level data ^[55]^ and DEqMS needs peptide or spectral counts ^[34]^. More descriptions about these normalization, imputation and DEA methods are presented in our **Supplementary results**. In summary, for DDA data, based on quantification results from FragPipe or maxquant, 3004 workflows are comparable (see **Supplementary file 2** for detail workflow lists), and for DIA data quantified with DIA-NN, 4800 workflows are accessible (**Supplementary file 2**).

### Performance evaluation metrics

The prepared datasets were processed to produce corresponding expression matrices (**Figure 1**). These expression matrices were then analyzed by workflows combining a particular expression matrix type, normalization method, MVI algorithm and DEA tool. The Benjamini-Hochberg method ^[56]^ is used for FDR control. A compiled table of DEA outcomes including protein ids, log2fold change values, p-values, adjust p-values (q-values) and contrast names, were then analyzed to compare performance.

Two performance measures were used including partial area under receiver operator characteristic curves (pAUC) and classification metrics:

#### partial area under receiver operator characteristic curves (pAUC)

The receiver operator characteristic curve (ROC) is generated by plotting True Positive Rates (TPR) against False Positive Rates (FPR) under various thresholds to classify samples into two categories ^[25]^. The area under the ROC (AUC) is a widely used performance indicators to evaluate the power of a classifier in classification tasks. The partial AUC is proposed for restricting the evaluation of given ROC curves in the range of FPRs that are considered interesting for diagnostic purposes ^[25]^. In our performance evaluations, the confidence score of a given protein to be a true differentially expressed protein is calculated by 1-q-value, where q-value is its Benjamini-Hochberg adjusted p-value. The FPR is restricted to FPR*≤*0.01, FPR*≤*0.05 and FPR*≤*0.1 respectively to calculate three performance indicators i.e., pAUC(0.01), pAUC(0.05) and pAUC(0.1) for a stricter performance evaluation.

#### Classification metrics

The classification metrics normalized Matthew’s correlation coefficient (nMCC) ^[26]^ and geometric mean of specificity and recall (G-mean) ^[57]^ were used as additional evaluation indicators where nMCC and G-mean are calculated as follows.

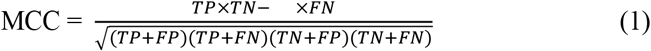

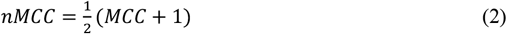

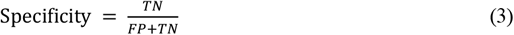

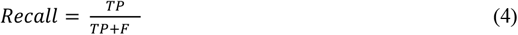

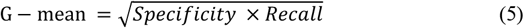

where True Positive (TP) is the truly differentially expressed protein has |logFC| *≥* 1 and q-value<0.05, False Positive (FP) is the truly non-differentially expressed protein has |logFC|*≥*1 and q-value<0.05, True Negative (TN) is the truly non-differentially expressed protein that can’t pass the threshold of |logFC|*≥*1 and q-value<0.05 while False Negative (FN) is the truly differentially expressed protein that cannot pass the threshold of |logFC|*≥*1 and q-value<0.05.The q-value is FDR adjusted p-value.

#### Ranking of workflows

Workflows were ranked by 5 performance metrics including pAUC(0.01), pAUC(0.05), pAUC(0.1), nMCC and G-mean separately. A workflow’s final rank is given by averaging its five ranks as shown below.

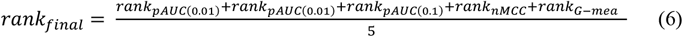

### Evaluation of benchmarking for workflow recommendation with leave-one-project-out cross-validation

To confirm whether our benchmarking results can be used for recommending optimal workflows for newcoming datasets, we conduct the leave-one-project-out cross-validation (LOPOCV). Taking the FragPipe data as an example, there are 4 projects data for benchmarking including yeast2099 (10 contrasts), yeast1819 (36 contrasts), CPTAC (with 4 sub-datasets, 40 contrasts) and human_ecoli (we generated 34 sub-datasets, 34 contrasts), see **Table 1** for more details. In a LOPOCV, each time, we use 3 out of 4 projects data to rank workflows (benchmarking), the contrasts in the remaining project data are regarded as newcoming datasets. We calculated the spearman correlation coefficient between the workflow ranks based on the benchmarking with the 3-project data and the true workflow ranks of the newcoming data. The higher the correlation is, the more accurate recommendations could be made with our benchmarking.

### Comparisons among choices in a single step of a workflow

After obtaining quantification results from a quantification platform, such as FragPipe, a comprehensive DEA workflow integrates several key selection steps, including:

a. An expression matrix that contains the expression levels of identified proteins;
b. A normalization method to reduce bias or noise;
c. An algorithm for imputing missing values in the selected expression matrix;
d. A DEA tool for conducting the final differential expression analysis.

Each step plays a crucial role. To examine the impact of a particular step, we simply maintained the options for other steps while varying the options of the step under investigation. To compare any two options for a given step, e.g., protein peak intensity and protein MaxLFQ intensity in step **a**, we calculated performance differences (mean pAUC(0.01) was used) of workflow pairs where they are alike in every other way except the choice of option. Different options in each step can be ranked by their pairwise comparisons. For a given step, we first count the frequencies of the options winning in pairwise mean performance comparisons. Then, the option with bigger frequency will be ranked higher. If two options have the same win frequencies, then their median performances will be compared for ranking.

### Ensemble Inference

Two ensemble strategies were used to verify our speculation including integration all (or 2 out of the 3) top 1^st^ workflows using spectral counts, protein intensities and peptide intensities (we name it ens_3inp or ens_2inp), and integration top K workflows in our overall rankings (ens_TK). Usually, the log2 fold change (log2FC) and FDR adjust p-value (q-value) are two key statistics to infer whether a protein is differentially expressed, our ensemble inference should integrate sub-workflows’ log2FCs and q-values into a log2FC and a q-value as ensembled statistics for the visited protein. For the integration of log2FC, we choose the log2FC having the biggest absolute value among all the sub-log2FCs as the ensembled log2FC. For q-values, five methods were used.

#### Hurdle model

The first one is used in Goeminne et al.’s hurdle model ^[38]^ where the p-values were transformed to z-values and combine them in a χ2 statistic:

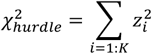

where χ2 statistic follows a χ2 distribution with t degrees of freedom if t out of K sub-p-values exist, and if t=1, then the integrated p-value equals to the existing one. Q-values will be obtained by FDR-adjustment with Benjamini-Hochberg’s method.

#### Fisher’s method

The second method is the Fisher’s combined probability test (Fisher’s method) ^[58]^. Fisher’s method firstly combines K sub-p-values as a χ2 statistic in the way of:

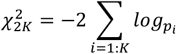

where *p*_*i*_ is the *i*th p-value, and χ2 has a chi-squared distribution with 2K degrees of freedom, then integrated p-value can be determined. Similarly, Benjamini-Hochberg adjusted q-values will be calculated.

#### Voting methods

The remaining three methods are based on voting strategies where we regard the K statistical tests as K voters. Anyone of the K p-values pass the threshold e.g., p-value < 0.05, then say the integrated test is significant, so we use minimum p-value (min p) of these K p-values as the integrated p-value. Similarly, if we adopt a most strict condition where we request all K p-values should pass, then maximum p-value (max p) will be used as integrated p-value. The last one is majority win, where if more p-values pass then integrated p-value pass, thus median of K p-values (median p) is used.

## Data availability

The raw proteomics data used in this work can be downloaded with the ProteomeXchange ID or Proteomic Data Commons Study Identifier listed in our **Table 1**. All the quantification results and our benchmarking results are available at our website: http://www.ai4pro.tech:3838/OpDEA_test/. Source data for all figures are provided with this paper.

## Code availability

The python and R codes used for benchmarking and data analysis are available at https://github.com/PennHui2016/OpDEA/tree/master/source_codes_for_benchmarking. The R package and its source codes of our tool OpDEA is available at: https://github.com/PennHui2016/OpDEA.

## Acknowledgement

This research/project is supported by the National Research Foundation, Singapore under its Industry Alignment Fund – Prepositioning (IAF-PP) Funding Initiative. Any opinions, findings and conclusions or recommendations expressed in this material are those of the author(s) and do not reflect the views of National Research Foundation, Singapore.

WWBG also acknowledges support from an MOE Tier 1 award (RS08/21).

## Author contributions

H.P. conceived and designed the experiments. H.P. collected the data, conducted the benchmarking, analyzed the data and deployed the webserver. H.P. and WWB.G. wrote the manuscript. H.W., W.K. and J.L. help improve the manuscript. J.L. and WWB.G supervised the study and acquired funding. All authors reviewed the manuscript and approved the manuscript.

## Additional information

Supplementary information

**Supplementary result**. Methods and Results not described in our main text.

**Supplementary file 1**. LOPOCV results for measuring the generalizability of our benchmarking.

**Supplementary file 2**. The workflow lists and our benchmarking results.

**Supplementary file 3**. Workflow performance levels and classification results with CatBoost.

**Supplementary file 4**. Frequent pattern mining results.

**Supplementary file 5**. Correlations between different metric-based workflow rankings.

**Supplementary file 6**. Detailed comparisons between options in each workflow step.

**Supplementary file 7**. Ensemble inference results.

## Reference

[1] Meissner, F., Geddes-McAlister, J., Mann, M., & Bantscheff, M. The emerging role of mass spectrometry-based proteomics in drug discovery. Nature Reviews Drug Discovery 21(9), 637–654 (2022).

[2] Niu, L., Thiele, M., Geyer, P. E., Rasmussen, D. N., Webel, H. E., Santos, A., Gupta, R., Meier, F., Strauss, M., Kjaergaard, M., et al. Noninvasive proteomic biomarkers for alcohol-related liver disease. Nature Medicine 28(6), 1277–1287 (2022).

[3] Langley, S. R. & Mayr, M. Comparative analysis of statistical methods used for detecting differential expression in label-free mass spectrometry proteomics. Journal of proteomics 129, 83–92 (2015).

[4] Ramus, C., Hovasse, A., Marcellin, M., Hesse, A.-M., Mouton-Barbosa, E., Bouyssié, D., Vaca, S., Carapito, C., Chaoui, K., Bruley, C., et al. Benchmarking quantitative label-free LC--MS data processing workflows using a complex spiked proteomic standard dataset. Journal of proteomics 132, 51–62 (2016).

[5] Välikangas, T., Suomi, T., & Elo, L. L. A comprehensive evaluation of popular proteomics software workflows for label-free proteome quantification and imputation. Briefings in bioinformatics 19(6), 1344–1355 (2018).

[6] Suomi, T., Seyednasrollah, F., Jaakkola, M. K., Faux, T., & Elo, L. L. ROTS: An R package for reproducibility-optimized statistical testing. PLoS computational biology 13(5), e1005562 (2017).

[7] Fröhlich, K., Brombacher, E., Fahrner, M., Vogele, D., Kook, L., Pinter, N., Bronsert, P., Timme-Bronsert, S., Schmidt, A., Bärenfaller, K., et al. Benchmarking of analysis strategies for data-independent acquisition proteomics using a large-scale dataset comprising inter-patient heterogeneity. Nature Communications 13(1), 2622 (2022).

[8] Lin, M.-H., Wu, P.-S., Wong, T.-H., Lin, I.-Y., Lin, J., Cox, J., & Yu, S.-H. Benchmarking differential expression, imputation and quantification methods for proteomics data. Briefings in Bioinformatics 23(3), bbac138 (2022).

[9] Sticker, A., Goeminne, L., Martens, L., & Clement, L. Robust summarization and inference in proteome-wide label-free quantification. Molecular & Cellular Proteomics 19(7), 1209–1219 (2020).

[10] Dowell, J. A., Wright, L. J., Armstrong, E. A., & Denu, J. M. Benchmarking quantitative performance in label-free proteomics. ACS omega 6(4), 2494–2504 (2021).

[11] Verhoeven, K. J., Simonsen, K. L., & McIntyre, L. M. Implementing false discovery rate control: increasing your power. Oikos 108(3), 643–647 (2005).

[12] Suomi, T. & Elo, L. L. Enhanced differential expression statistics for data-independent acquisition proteomics. Scientific reports 7(1), 5869 (2017).

[13] Kong, A. T., Leprevost, F. V., Avtonomov, D. M., Mellacheruvu, D., & Nesvizhskii, A. I. MSFragger: ultrafast and comprehensive peptide identification in mass spectrometry--based proteomics. Nature methods 14(5), 513–520 (2017).

[14] Demichev, V., Messner, C. B., Vernardis, S. I., Lilley, K. S., & Ralser, M. DIA-NN: neural networks and interference correction enable deep proteome coverage in high throughput. Nature methods 17(1), 41–44 (2020).

[15] Huber, W., Von Heydebreck, A., Sültmann, H., Poustka, A., & Vingron, M. Variance stabilization applied to microarray data calibration and to the quantification of differential expression. Bioinformatics 18(suppl_1), S96–S104 (2002).

[16] Stekhoven, D. J. & Bühlmann, P. MissForest—non-parametric missing value imputation for mixed-type data. Bioinformatics 28(1), 112–118 (2012).

[17] Kerr, M. K., Martin, M., & Churchill, G. A. Analysis of variance for gene expression microarray data. Journal of computational biology 7(6), 819–837 (2000).

[18] Tusher, V. G., Tibshirani, R., & Chu, G. Significance analysis of microarrays applied to the ionizing radiation response. Proceedings of the National Academy of Sciences 98(9), 5116–5121 (2001).

[19] Hatfield, G. W., Hung, S.-p., & Baldi, P. Differential analysis of DNA microarray gene expression data. Molecular microbiology 47(4), 871–877 (2003).

[20] Tyanova, S., Temu, T., & Cox, J. The MaxQuant computational platform for mass spectrometry-based shotgun proteomics. Nature protocols 11(12), 2301–2319 (2016).

[21] Espndola, R. P. & Ebecken, N. F. On extending f-measure and g-mean metrics to multi-class problems. WIT Transactions on Information and Communication Technologies 35, 25–34 (2005).

[22] Pursiheimo, A., Vehmas, A. P., Afzal, S., Suomi, T., Chand, T., Strauss, L., Poutanen, M., Rokka, A., Corthals, G. L., & Elo, L. L. Optimization of statistical methods impact on quantitative proteomics data. Journal of proteome research 14(10), 4118–4126 (2015).

[23] Paulovich, A. G., Billheimer, D., Ham, A.-J. L., Vega-Montoto, L., Rudnick, P. A., Tabb, D. L., Wang, P., Blackman, R. K., Bunk, D. M., Cardasis, H. L., et al. Interlaboratory study characterizing a yeast performance standard for benchmarking LC-MS platform performance. Molecular & Cellular Proteomics 9(2), 242–254 (2010).

[24] Gotti, C., Roux-Dalvai, F., Joly-Beauparlant, C., Mangnier, L., Leclercq, M., & Droit, A. DIA proteomics data from a UPS1-spiked E. coli protein mixture processed with six software tools. Data in Brief 41, 107829 (2022).

[25] McClish, D. K. Analyzing a portion of the ROC curve. Medical decision making 9(3), 190–195 (1989).

[26] Chicco, D. & Jurman, G. The advantages of the Matthews correlation coefficient (MCC) over F1 score and accuracy in binary classification evaluation. BMC genomics 21, 1–13 (2020).

[27] Cox, J., Hein, M. Y., Luber, C. A., Paron, I., Nagaraj, N., & Mann, M. Accurate proteome-wide label-free quantification by delayed normalization and maximal peptide ratio extraction, termed MaxLFQ. Molecular & cellular proteomics 13(9), 2513–2526 (2014).

[28] Karpievitch, Y., Stanley, J., Taverner, T., Huang, J., Adkins, J. N., Ansong, C., Heffron, F., Metz, T. O., Qian, W.-J., Yoon, H., et al. A statistical framework for protein quantitation in bottom-up MS-based proteomics. Bioinformatics 25(16), 2028–2034 (2009).

[29] Wei, R., Wang, J., Su, M., Jia, E., Chen, S., Chen, T., & Ni, Y. Missing value imputation approach for mass spectrometry-based metabolomics data. Scientific reports 8(1), 1–10 (2018).

[30] Prokhorenkova, L., Gusev, G., Vorobev, A., Dorogush, A. V., & Gulin, A. CatBoost: unbiased boosting with categorical features. Advances in neural information processing systems 31 (2018).

[31] Han, J., Pei, J., Yin, Y., & Mao, R. Mining frequent patterns without candidate generation: A frequent-pattern tree approach. Data mining and knowledge discovery 8, 53–87 (2004).

[32] Smyth, G. K. Limma: linear models for microarray data. Bioinformatics and computational biology solutions using R and Bioconductor, 397–420 (2005).

[33] van Ooijen, M. P., Jong, V. L., Eijkemans, M. J., Heck, A. J., Andeweg, A. C., Binai, N. A., & van den Ham, H.-J. Identification of differentially expressed peptides in high-throughput proteomics data. Briefings in bioinformatics 19(5), 971–981 (2018).

[34] Zhu, Y., Orre, L. M., Tran, Y. Z., Mermelekas, G., Johansson, H. J., Malyutina, A., Anders, S., & Lehtiö, J. DEqMS: a method for accurate variance estimation in differential protein expression analysis. Molecular & Cellular Proteomics 19(6), 1047–1057 (2020).

[35] Kong, W., Hui, H. W. H., Peng, H., & Goh, W. W. B. Dealing with missing values in proteomics data. Proteomics 22(23-24), 2200092 (2022).

[36] Audigier, V., Husson, F., & Josse, J. Multiple imputation for continuous variables using a Bayesian principal component analysis. Journal of statistical computation and simulation 86(11), 2140–2156 (2016).

[37] Goeminne, L. E., Gevaert, K., & Clement, L. Peptide-level robust ridge regression improves estimation, sensitivity, and specificity in data-dependent quantitative label-free shotgun proteomics. Molecular & Cellular Proteomics 15(2), 657–668 (2016).

[38] Goeminne, L. J., Sticker, A., Martens, L., Gevaert, K., & Clement, L. MSqRob takes the missing hurdle: uniting intensity-and count-based proteomics. Analytical chemistry 92(9), 6278–6287 (2020).

[39] Schwender, H. & Ickstadt, K. Empirical Bayes analysis of single nucleotide polymorphisms. BMC bioinformatics 9(1), 1–15 (2008).

[40] Crookston, N. L. & Finley, A. O. yaImpute: an R package for kNN imputation. Journal of Statistical Software 23, 1–16 (2008).

[41] UniProt: the Universal Protein knowledgebase in 2023. Nucleic Acids Research 51(D1), D523–D531 (2023).

[42] Käll, L., Canterbury, J. D., Weston, J., Noble, W. S., & MacCoss, M. J. Semi-supervised learning for peptide identification from shotgun proteomics datasets. Nature methods 4(11), 923–925 (2007).

[43] Nesvizhskii, A. I., Keller, A., Kolker, E., & Aebersold, R. A statistical model for identifying proteins by tandem mass spectrometry. Analytical chemistry 75(17), 4646–4658 (2003).

[44] Yu, F., Haynes, S. E., & Nesvizhskii, A. I. IonQuant enables accurate and sensitive label-free quantification with FDR-controlled match-between-runs. Molecular & Cellular Proteomics 20 (2021).

[45] Yu, S.-H., Kyriakidou, P., & Cox, J. Isobaric matching between runs and novel PSM-level normalization in MaxQuant strongly improve reporter ion-based quantification. Journal of proteome research 19(10), 3945–3954 (2020).

[46] Cox, J., Neuhauser, N., Michalski, A., Scheltema, R. A., Olsen, J. V., & Mann, M. Andromeda: a peptide search engine integrated into the MaxQuant environment. Journal of proteome research 10(4), 1794–1805 (2011).

[47] Gatto, L. & Lilley, K. S. MSnbase-an R/Bioconductor package for isobaric tagged mass spectrometry data visualization, processing and quantitation. Bioinformatics 28(2), 288–289 (2012).

[48] White, I. R., Royston, P., & Wood, A. M. Multiple imputation using chained equations: issues and guidance for practice. Statistics in medicine 30(4), 377–399 (2011).

[49] Gentleman, R. C., Carey, V. J., Bates, D. M., Bolstad, B., Dettling, M., Dudoit, S., Ellis, B., Gautier, L., Ge, Y., Gentry, J., et al. Bioconductor: open software development for computational biology and bioinformatics. Genome biology 5(10), 1–16 (2004).

[50] Zhang, X., Smits, A. H., Van Tilburg, G. B., Ovaa, H., Huber, W., & Vermeulen, M. Proteome-wide identification of ubiquitin interactions using UbIA-MS. Nature protocols 13(3), 530–550 (2018).

[51] Pavelka, N., Pelizzola, M., Vizzardelli, C., Capozzoli, M., Splendiani, A., Granucci, F., & Ricciardi-Castagnoli, P. A power law global error model for the identification of differentially expressed genes in microarray data. BMC bioinformatics 5(1), 1–12 (2004).

[52] Ahlmann-Eltze, C. and Anders, S. proDA: probabilistic dropout analysis for identifying differentially abundant proteins in label-free mass spectrometry. Biorxiv, 661496 (2019).

[53] Robinson, M. D., McCarthy, D. J., & Smyth, G. K. edgeR: a Bioconductor package for differential expression analysis of digital gene expression data. Bioinformatics 26(1), 139–140 (2010).

[54] Baggerly, K. A., Deng, L., Morris, J. S., & Aldaz, C. M. Differential expression in SAGE: accounting for normal between-library variation. Bioinformatics 19(12), 1477–1483 (2003).

[55] Choi, M., Chang, C.-Y., Clough, T., Broudy, D., Killeen, T., MacLean, B., & Vitek, O. MSstats: an R package for statistical analysis of quantitative mass spectrometry-based proteomic experiments. Bioinformatics 30(17), 2524–2526 (2014).

[56] Ferreira, J. A. The Benjamini-Hochberg method in the case of discrete test statistics. The international journal of biostatistics 3(1) (2007).

[57] Xuan, X., Lo, D., Xia, X., & Tian, Y. Evaluating defect prediction approaches using a massive set of metrics: An empirical study. In Proceedings of the 30th Annual ACM Symposium on Applied Computing, 1644–1647, (2015).

[58] Elston, R. On Fisher’s method of combining p-values. Biometrical journal 33(3), 339–345 (1991).

